# Starvation improves epithelial fitness by selectively extruding DNA damaged cells

**DOI:** 10.64898/2026.09.01.748489

**Authors:** Lily A. Gates, Dustin C. Bagley, Konstantinos Kalyviotis, John Fadul, Tania Auchynnikava, Maria Voronkov, Jody Rosenblatt

## Abstract

During homeostasis, crowded cells with the lowest energy levels are eliminated by extrusion via Piezo1 signalling to maintain constant cell numbers. However, crowding-induced extrusion does not necessarily remove damaged or otherwise unfit cells. Here, we show that glucose or glutamine starvation triggers a rapid, regulated wave of extrusion, called starvation-induced cell extrusion (STICE), that selectively eliminates cells bearing DNA damage markers via a p53-dependent, Piezo1-independent pathway, improving monolayer fitness. Unlike non-extruding cells, which recycle contents through autophagy and lysosomal digestion, p53-activated cells instead use LC3 to drive lysosomal exocytosis, promoting extrusion signalling. By eliminating defective and transformed cells, STICE confers resistance to damage and apoptotic stimuli in the remaining monolayer. STICE thus acts as a tissue-level analogue of autophagy: rather than improving individual cells by digesting and recycling damaged components, it improves tissue fitness by eliminating substandard cells.

## Main

Epithelial cells work tightly together to build protective barriers and are necessary for proper organ function. As the first line of defence, epithelia must withstand harsh environments and continuously renew through cell division and death^1,2^. To match the numbers of cells dying to those dividing while keeping a tight barrier, epithelia seamlessly extrude excess or unwanted cells^1,3^. A cell destined to extrude emits sphingosine-1-phosphate (S1P), which binds the G-protein coupled receptor, S1P2, triggering basolateral actomyosin contraction that squeezes the cell out apically from the layer, preserving barrier function^4,5^. Cell extrusion is a primordial process conserved from sea sponge^6^ to humans^1^ that drives epithelial cell death and is essential for preventing inflammation and neoplastic growth^7,8^. Most extrusions are live, resulting from overcrowding, with cells subsequently dying from a lack of survival signalling^3^, however, numerous other triggers can activate extrusion in different contexts. Apoptotic cells are extruded early to prevent barrier breaches and to eliminate topological defects within the epithelial fabric^9,10^. Moreover, live cells with replicative stress or that are otherwise unfit, can be targeted for extrusion via p53 activation^11,12^. Additionally, extrusion is emerging as a fundamental step in innate immunity, potentially conserved back to epithelioid organisms that lack a blood-based immune system^13^.

While many triggers elicit extrusion^10–13^, the most common route is crowding-induced live cell extrusion^1^, which mechanically matches the number of cells extruding to those dividing at rates as high as ∼10 billion cells/hour in the gut epithelium alone^14^. While crowding activates extrusion via the stretch-activated channel Piezo1, the crowded cells targeted for extrusion have proportionally less ATP^15^. Given that energetically weaker cells are selected for extrusion, here, we investigated if nutrient starvation would trigger excess extrusion of suboptimal cells.

## Results

To test how epithelia respond to starvation, we treated two epithelial cell lines (MDCKTII and 16HBE) with Hank’s Balanced Salt Solution (HBSS) versus complete media (fed) and imaged by phase-contrast live over four hours. HBSS significantly increased extrusion within as little as one hour (**Fig. 1A-B, Extended Data Fig. 1, Supplementary Video 1-2**). Changing to complete medium did not have the same response, ruling out shear stress as driving this response. Strikingly, starvation did not cause ongoing extrusion but was instead limited to a single wave peaking at two hours, returning to baseline at four hours, which we term Starvation-Induced Cell Extrusion (STICE) (**Fig. 1C**). This self-limiting wave of extrusion may protect epithelial numbers, as ongoing extrusion would damage the barrier, causing inflammation.

**Fig 1.**
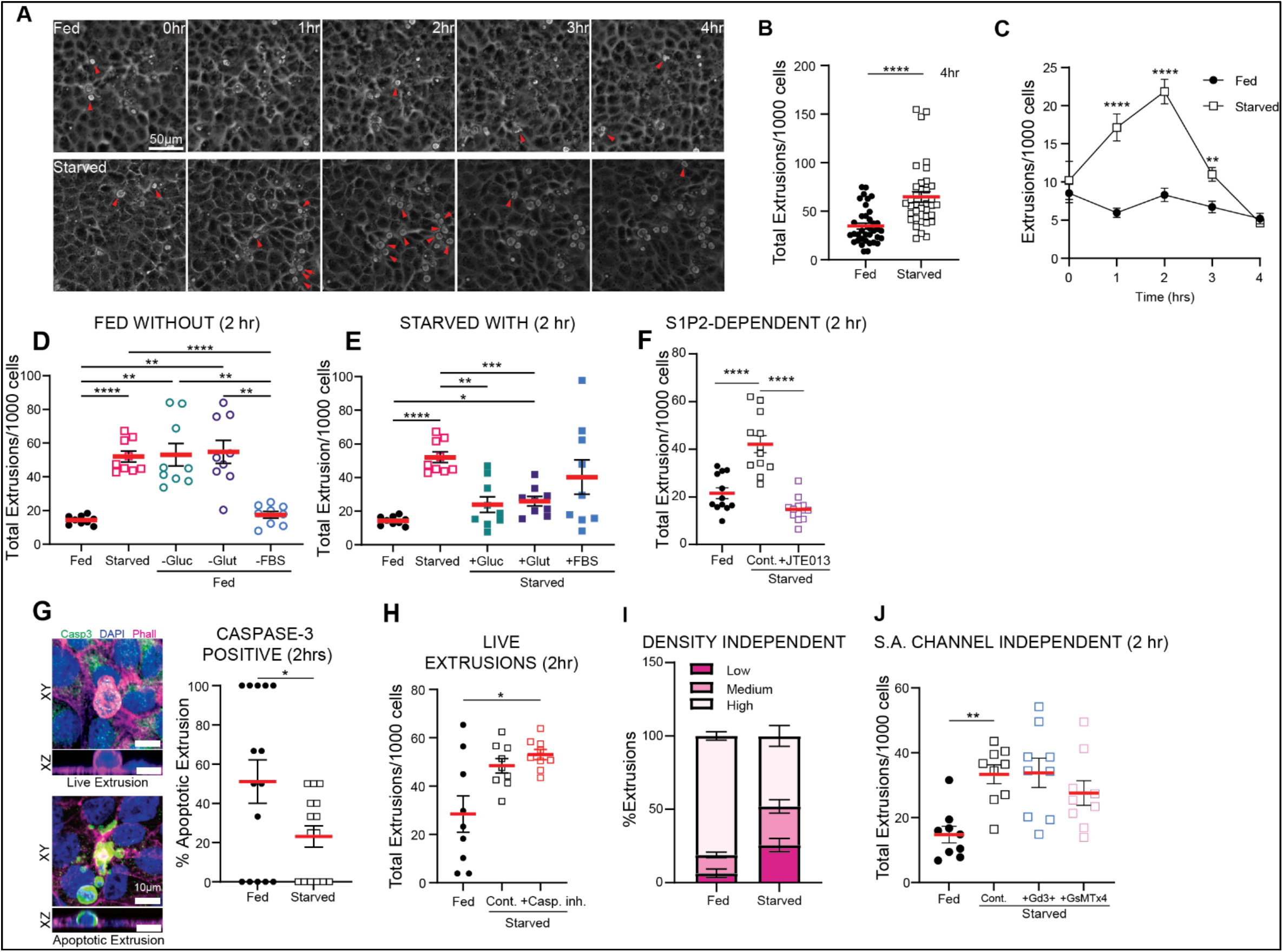
Starvation induces a wave of live cell extrusion (STICE) **A**, Stills from hourly intervals of phase-contrast live imaging of MDCKII monolayers cultured in normal growth medium (Fed, top) or HBSS (Starved, bottom), imaged every 10 min for 4 h. Red arrows indicate extrusions. Scale bar, 50 µm. **B**, Normalised extrusions per 1,000 cells over 4 h time course. n=36 fields of view (12 biological replicates); p****≤0.0001 from Mann-Whitney t-test. **C**, Time course of total extrusions per 1,000 cells scored hourly. n=36 fields of view (12 biological replicates). **D-E**, Cumulative extrusion counts per 1,000 cells after 2 h in Fed or Starved, with glucose, glutamine or FBS withdrawn from Fed medium (**D**) or supplemented into Starved medium (**E**). n=9 fields of view (3 biological replicates); p*≤0.05, p**≤0.01, p***≤0.001, p****≤0.0001, from Welch’s one-way ANOVA with Dunnett’s multiple-comparisons test. **F**, Cumulative extrusion counts per 1,000 cells after 2 h in Fed or Starved ± 10 µM JTE013. N=12 fields of view (4 biological replicates); p****≤0.0001 from Welch’s one-way ANOVA, with Dunnett’s multiple-comparisons test. **G**, Representative confocal projection images of non-apoptotic (active caspase-3 negative, top) and apoptotic (active-caspase-3 positive, bottom) extrusions from confluent monolayers fixed after 2 h in Fed or Starved with XZ projections below. Scale bar, 10 µm. Cumulative percentage of caspase-3-positive extrusions over 2 h (right). n=15 fields of view (3 biological replicates); p*≤0.05 from Welch’s t-test. **H**, Total extrusions per 1,000 cells after 2 h in Fed and Starved, with or without 20 µM Z-DEVD-FMK. n=9 fields of view (3 biological replicates); p*≤0.05, from Welch’s one-way ANOVA, with Dunnett’s multiple comparisons test. **I**, Extrusions scored in regions of low, medium and high crowding, and expressed as percentages of total extrusions in Fed and Starved conditions. n=6 fields of view per density (5 biological replicates). **J**, Total extrusions per 1,000 cells after 2 h in Fed and Starved ± 10 µM GdCl_3_ or 50 µM GsMTx4. n=9 fields of view (3 biological replicates); p**≤0.01, from Kruskal-Wallis test with Dunn’s multiple-comparisons test.

To identify factors responsible for STICE, we removed or added components from complete medium. Complete medium lacking either glutamine or glucose induced the same percentage of extrusions as HBSS, while serum-free medium, lacking growth factors, had no effect (**Fig. 1D**). Conversely, supplementing HBSS with glucose or glutamine prevented STICE, whereas adding foetal bovine serum did not (**Fig.1E**). Together, these results show that glucose and glutamine regulate STICE, while serum plays no role.

We next tested if STICE requires canonical extrusion signalling pathways. Disrupting S1P signalling through the S1P2 receptor^4,5^ with the S1P2 antagonist JTE013, reduced STICE rates comparable to fed controls (**Fig. 1F**), indicating that STICE uses the same canonical S1P signalling required for other extrusion pathways^10–13^. Immunostaining for active caspase-3 revealed that most STICE events were live-cell extrusions (**Fig. 1G**). Caspase-3 inhibition with Z-DEVD-FMK did not reduce STICE rates, confirming that STICE does not require apoptotic signalling (**Fig. 1H**). Because most live-cell extrusion is normally driven by crowding, we next tested whether the Piezo1-dependent crowding pathway also governs STICE. Interestingly, however, STICE was not restricted to crowded regions of the monolayer, like fed controls (**Fig. 1I**). Further, inhibiting stretch-activated channels with gadolinium or GsMTx4^16^ had no impact on STICE (**Fig. 1J**). Together, these results indicate that Piezo1 and crowding do not regulate STICE, suggesting a distinct upstream trigger activates S1P-mediated extrusion in response to starvation or that a different population is targeted by starvation.

Having ruled out crowding as an upstream trigger, we next asked what marks cells for elimination by STICE. Immunostaining fed and starved monolayers for DNA damage markers, showed that γ-H2AX, a marker of double-strand DNA breaks^17^, was elevated in extruding cells following starvation, compared to fed conditions (**Fig. 2A**), as was active phospho-p53, a master regulator of DNA repair and apoptosis (**Fig. 2B**). To test if STICE targeted cells were functionally compromised, we replated extruded cells from fed and starved monolayers to assess their fitness (**Fig. 2C**). We found that far fewer STICE-derived cells survived after 24 hours (**Fig. 2D**), and those that did were less proliferative than those from fed monolayers, as indicated by reduced Ki67 staining^18^ (**Fig. 2E**). Thus, STICE eliminates less healthy cells than those from crowding-induced extrusion during homeostasis^1^. This selective elimination of less-fit cells may explain the self-limiting nature of the extrusion wave following starvation.

**Fig 2.**
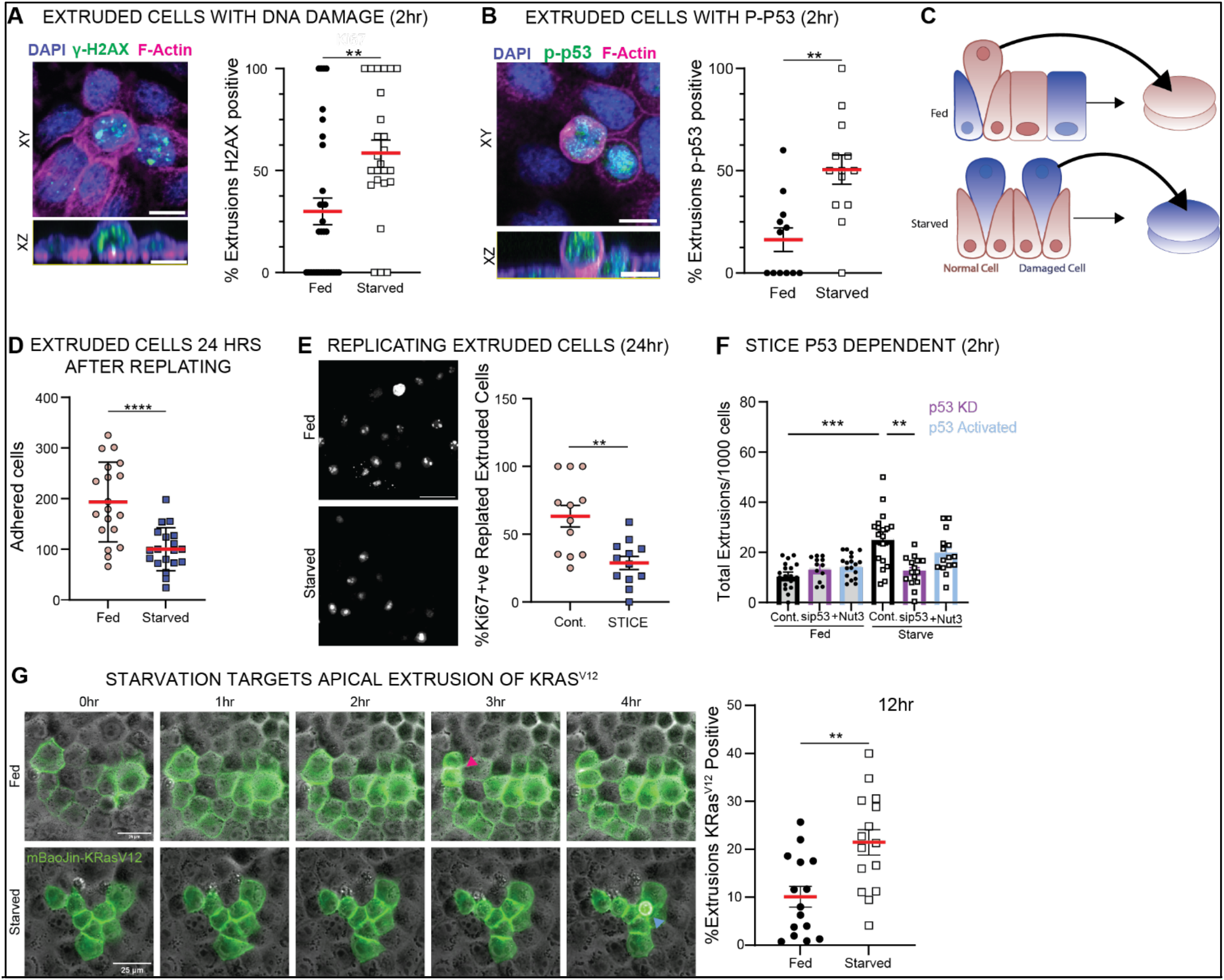
STICE selectively extrudes damaged and transformed cells via p53 activation. **A-B**, Representative confocal projection images of γ-H2AX-positive (**A**, left) or p-p53-positive (**B**, left) cells extruding from confluent MDCKT2 monolayers fixed after 2 h in DMEM (Fed) or HBSS (Starved), with XZ projections below. Scale bar, 10 µm. Percentage of γ-H2AX-positive (**A**) or p-p53-positive (**B**) extrusions at 2 h (right). n=12 fields of view (3 biological replicates) with one-way ANOVA (**A**; p**≤0.01) or Mann-Whitney test (**B**; p***≤0.001). **C**, Schematic of re-plating assay: confluent MDCKII monolayers Fed and Starved for 2 h, lightly trypsinized to isolate extruded cells, which were replated and grown for 24 h. **D**, Number of extruded cells adhered to the plate at 24 h from Fed and Starved extrusion. n=19 (5 biological replicates). p****≤0.0001 from an unpaired Welch’s t-test. **E**, Representative confocal images of replated extruded cells stained for Ki67 and percentage Ki67-positive cells in Fed and Starved conditions (right). n=12 (3 biological replicates); p**≤0.01 from an unpaired Welch’s t-test. **F**, Total extrusions per 1,000 cells after 2 h in p53-knockdown (sip53) and control (-) MDCKTII cells, in Fed or Starved ± 10 µM Nutlin-3 (Nut3). n=18 fields of view (6 biological replicates); p**≤0.01, p****≤0.0001 from Welch’s one-way ANOVA. **G**, Wild-type and doxycycline-inducible KRasV12 MDCKTII cells combined 50:1, induced 24 h prior imaging, incubated in fed or starved, and imaged for 12 h at 10 min intervals. Representative stills over the first 4 h (left); pink arrows, divisions; blue arrows, apical extrusions. Scale bar, 25 µm. Percentage of total extrusions over 12 h that were mBaoJin-KRasV12-positive in fed and starved (right). n=15 fields of view (5 biological replicates); p**≤0.01 from unpaired t-test.

Although STICE occurs independently of apoptosis, live-cell extrusion driven by mechanical cell competition or replicative stress requires p53^11,12^, activated in STICE cells (**Fig. 2B**). To investigate whether STICE requires p53, we tested if p53 knockdown reduced STICE. While p53 knockdown did not affect homeostatic extrusion in fed conditions, it significantly reduced STICE (**Fig. 2F**). However, activating p53 with Nutlin-3 did not further increase STICE rates under starvation (**Fig. 2F**), suggesting p53 activation and starvation are not sufficient to promote extrusion.

As many transformed cells contain DNA damage^19^, we next investigated if STICE could be co-opted to selectively remove transformed cells. Using MDCK monolayers mosaically expressing inducible KRas^V12^, a driver of mutation of many aggressive cancers^20^. We found that HBSS starvation selectively targeted KRas^V12^-transformed cells for STICE, which was not seen in complete medium (**Fig. 2G, Extended Data Fig. 2**). Thus, starvation appears to selectively target transformed cells as well as defective cells for extrusion.

To interrogate pathways downstream of glucose and glutamine starvation that might activate STICE, we investigated candidates in macroautophagy (hereafter, autophagy), which is similarly induced by starvation ^21–23^. Autophagy is an ancient cellular process conserved across eukaryotes that sequesters cytoplasmic components and damaged organelles in double-membrane vesicles for lysosomal degradation and recycling^21–23^ (**Fig. 3A**). Since mTOR is a master regulator of metabolism and inhibitor of autophagy in response to nutrient availability, we tested whether Torin1, a potent and selective mTOR inhibitor^24^ could induce extrusion under fed conditions (**Fig. 3B**). Torin1 increased extrusion, but not to the levels of glucose or glutamine starvation. Additionally, inhibiting the upstream autophagy kinase ULK1^25,26^ with MRT68921^27^ did not affect STICE (**Fig. 3B**), suggesting STICE uses a non-canonical form of autophagy or another pathway downstream of starvation. ATG5, which with ATG13 and ATG16L1, mediates LC3 lipidation required for autophagosome production^23,28^ in both canonical and non-canonical autophagy. We found that siATG5-mediated knockdown greatly reduced STICE without impacting homeostatic fed extrusion (**Fig. 3C**), suggesting that STICE requires autophagosomes but not through a canonical autophagy pathway.

**Fig 3.**
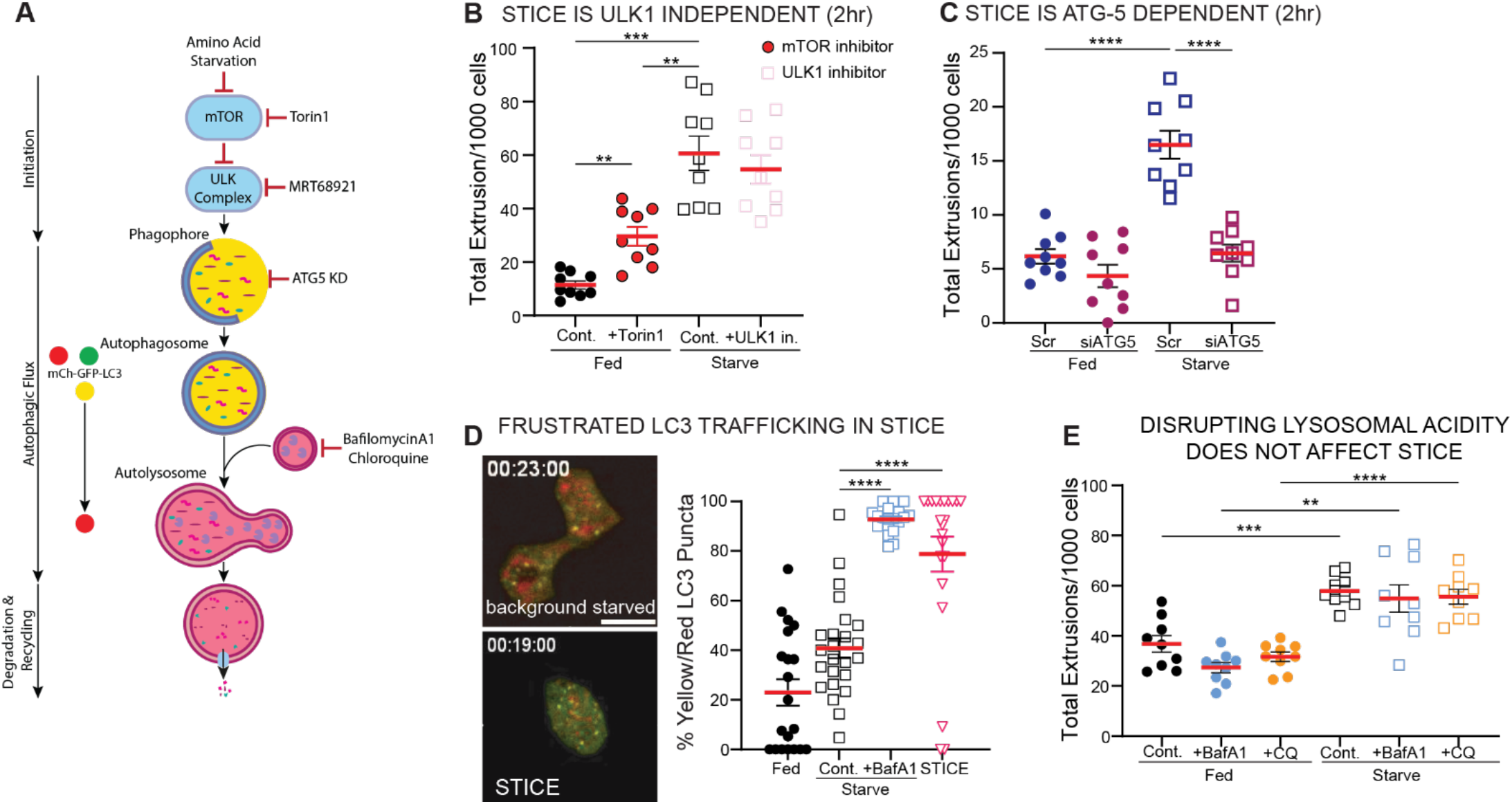
STICE requires ATG5 and LC3 but not canonical autophagy for extrusion. **A**, Simplified schematic of the autophagy pathway showing inhibitor targets and the mCherry-GFP-LC3 reporter, in which autophagosomes appear yellow and acidification upon lysosomal fusion quenches GFP, shifting puncta to red. **B**, Total extrusions per 1,000 cells after 2 h in DMEM (Fed) ± 1 µM Torin1, and HBSS (Starved) with or without 1 µM MRT68921 (ULK1 inhibitor). n=9 fields of view (3 biological replicates); p**≤0.01, p***≤0.001 from Welch’s one-way ANOVA with Dunnett’s T3 multiple-comparisons test. **C**, Total extrusions per 1,000 cells after 2 h in ATG5 knockdown (siATG5) and control (Src) MDCKTII cells, in Fed and Starved. n=9 fields of view (3 biological replicates); p****≤0.0001 from Welch’s one-way ANOVA, with Dunnett’s T3 multiple-comparisons. **D**, Representative live confocal images of MDCKT-mCherry-GFP-LC3 monolayers starved for 2 h, showing a background monolayer cell (top), and a pre-extruding STICE cell 5 min before extrusion (bottom). Scale bar, 10 µm. Percentage of yellow versus red puncta quantified in background monolayer cells (fed, starved, and starved with 100 nM bafilomycin A1) and in pre-extruding STICE cells. n=3 biological replicates; p****≤0.0001 from Kruskal-Wallis test with Dunn’s multiple-comparisons test. **E**, Total extrusions per 1,000 cells after 2 h in Fed and Starved ± 100 nM bafilomycinA1 or 50 µM chloroquine. n=9 fields of view (3 biological replicates); p**≤0.01, p***≤0.001, p****≤0.0001 from Welch’s one-way ANOVA with Dunnett’s T3 multiple-comparisons test.

A key step in canonical autophagy is autophagosome fusion with lysosomes for degradation to recycling cellular components^29,30^. To test whether STICE requires this autophagosome fusion to lysosomes, or autophagic flux, we used the mCherry-GFP-LC3 reporter, where autophagosomes appear yellow (green and red), while those fused to lysosomes appear red^31,32^ (**Fig. 3A**). While LC3 puncta in background starved cells shifted from yellow to red, those in cells undergoing STICE remained yellow (**Fig. 3D, Supplementary Video 3-4**), suggesting that autophagosomes in cells undergoing STICE do not fuse to lysosomes or that lysosomes are deacidified. Additionally, chloroquine and bafilomycinA1, which reduce lysosome acidity^33,34^, did not affect STICE rates (**Fig. 3E**), further supporting a model in which lysosomal degradation via canonical autophagy is dispensable for STICE. However, neither chloroquine nor bafilomycinA1 in fed conditions induced extrusion, suggesting deacidification of the lysosomes is not sufficient to induce STICE.

To assess lysosomal function, we used LysoTracker, a dye that fluoresces only in acidic compartments. Cells both before and during STICE had reduced LysoTracker puncta relative to starved and fed monolayer cells (**Fig. 4A**), suggesting that their lysosomes were either reduced in number or unable to maintain a low pH. Because p53 can promote lysosomal membrane permeabilization (LMP) via activation and translocation of truncated BID (tBID) to lysosomal membranes, causing membrane disruption and leakage of lysosomal enzymes^35^, we tested whether reduced LysoTracker signal reflected leaky lysosomes. RFP-Galectin-3, which forms puncta specifically in cells with permeabilised lysosomes^36^, showed little to no accumulation in STICE cells prior to extrusion (**Fig. 4B**). By contrast, permeabilising lysosomes with L-leucyl-L-leucine methyl ester (LLOMe), caused significant puncta formation^35^ (**Fig. 4B**).

**Fig 4:**
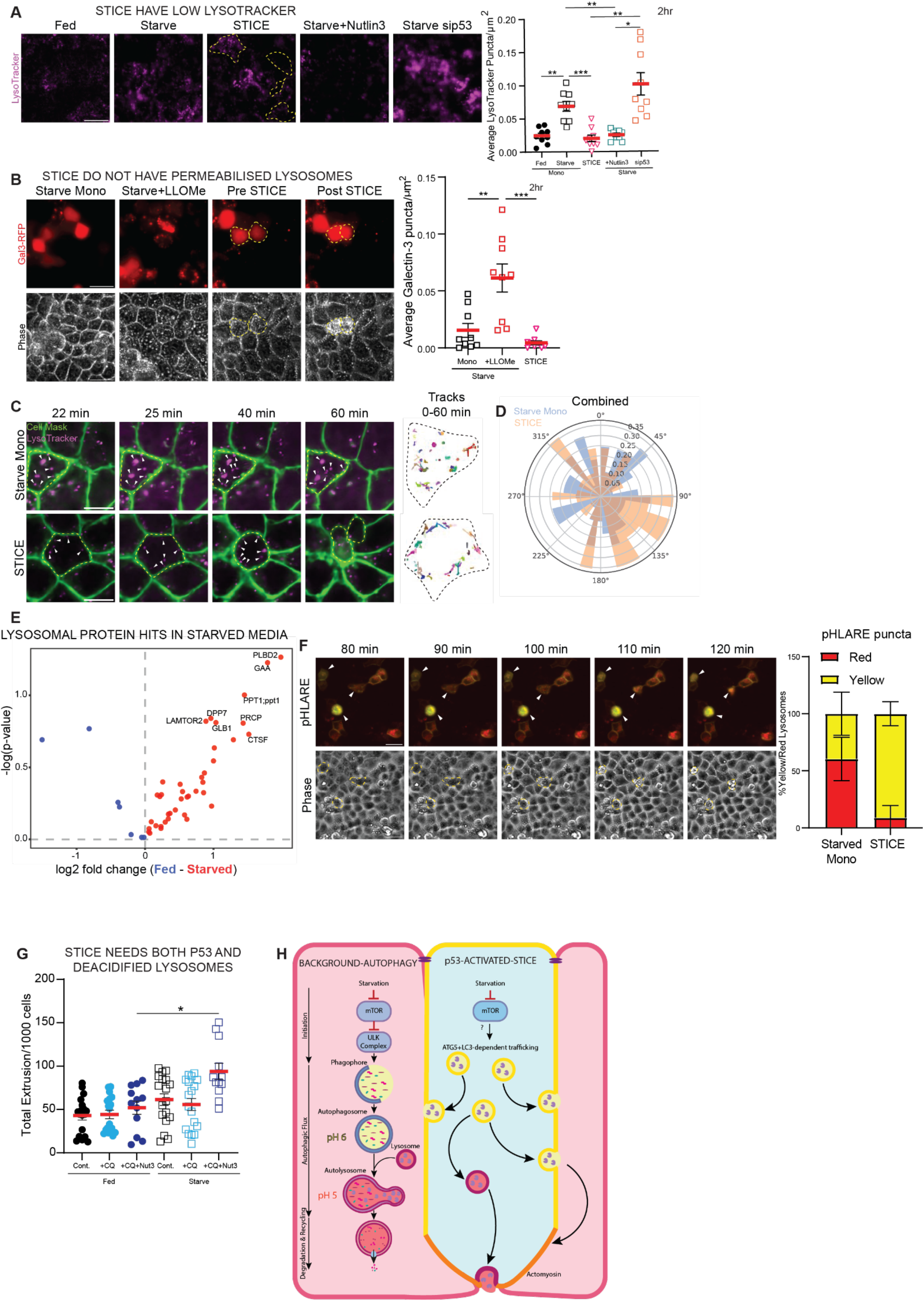
STICE traffics lysosomes to plasma membrane. **A**, Representative live confocal images of MDCKTII cells in DMEM (fed), HBSS (starve) for 2h ± 10 µM Nutlin-3, or with p53 knockdown (sip53), stained with LysoTracker. Pre-extruding cells are outlined in yellow dashed lines. Scale bar 25 µm. LysoTracker puncta from these treatments is shown to the right. n=9 averaged fields of view (3 biological replicates); p*≤0.05, p**≤0.01, p***≤0.001 from one-way Welch’s ANOVA with Dunnett’s T3 multiple-comparisons test. **B**, MDCKTII cells expressing Galectin-3-RFP were starved ± 1 mM LLOMe, imaged live for 4 h at 10 min intervals. Representative stills of Gal3-RFP-positive extrusion (top) with corresponding phase images (bottom) over 2 h. Extruding cell outlines by yellow dashed lines. Scale bar 25 µm Galectin-3 puncta per µm^2^ in starved monolayer ± LLOMe and in STICE cells at 2 h (right). n=9 fields of view (3 biological replicates); p*≤0.05, p***≤0.001 from Kruskal-Wallis test with Dunn’s multiple comparisons test. **C**, Representative live confocal images of MDCKTII cells starved for 2h, stained with LysoTracker and CellMask, images every 15s. Bottom panel highlights an extruding cell. White arrows correspond to LysoTracker puncta. Cell used for 60 min LysoTracker tracking image (left), is highlighted in yellow. Scale bar 10 µm. Tracks were generated with TrackMate. **D**, Combined rose plot of tracking data from LysoTracker in both starved monolayers and starved extruding cells. n=11 STICE cells, n=21 starved monolayer cells. **E**, Volcano plot of lysosomal protein hits identified in the media secretome of fed (blue) and starved (red) monolayers at 2 h, labelling the top 8 hits in starved. **F**, Representative stills of MDCKTII cells expressing pHLARE (top), with corresponding phase images (bottom), starved for 4 h at 10 min intervals. Extruding cells are highlighted with white arrows (top) and yellow dashed outline (bottom). Scale bar 25 µm. Quantification of ratio of yellow:red lysosomal pHLARE puncta from starved monolayer and STICE cells across 4 h (right). **G**, Total extrusions per 1,000 cells after 2 h in Fed and Starved ± 50 µM chloroquine (CQ) and 10 µM Nutlin-3. n=12 fields of view (4 biological replicates); p*≤0.05 from Kruskal-Wallis test with Dunn’s multiple-comparisons test. **H**, Proposed model; starvation induces autophagy in most cells but STICE in cells with DNA-damage but activation exocytosis of acidic and de-acidified lysosomes, secreting extrusion signals and enzymes that degrade cell-substrate and cell-cell junctions.

Since lysosomal permeabilization does not account for the reduced LysoTracker signal, we instead tracked lysosomes directly at high resolution and high speed. Notably, there is more peripheral lysosomal movement in STICE cells, with more central confined lysosomal movement in starved non-extruding cells (**Fig. 4C-D, Supplementary Videos 5-6**). Similar results were noted with LAMP1-tdTomato at high temporal and spatial resolution, also showing membrane buckling (**Supplementary Videos 7-8**), consistent with STICE cells exhibiting increased lysosomal exocytosis^37^. Secretomics of conditioned medium from HBSS-starved versus fed monolayers indicated that lysosomal proteins, including PLBD2, GAA and GLB1, were preferentially secreted from starved monolayers (**Fig. 4E**), further suggesting that lysosomes in cells undergoing STICE release their contents via exocytosis. Together, these findings may suggest a model in which STICE cells use ATG5 and LC3 to traffic lysosomes to the plasma membrane to signal extrusion.

However, because lysosomes typically degrade most proteins owing to their acidic, hydrolase rich content, they seem an unlikely source of extrusion-promoting factors. To test whether lysosomes in cells undergoing STICE have low pH, we used pH Lysosomal Activity Reporter (pHLARE), a genetically encoded LAMP1 construct tagged with both mCherry and sfGFP that allows acidified (red) and non-acidified (yellow) lysosomes to be distinguished^38^. We found that cells undergoing STICE had predominantly yellow, non-acidified lysosomes, consistent with the LC3 flux data in **Fig. 3D**, compared with background starved cells (**Fig. 4F, Supplementary Video 9**).

To test whether p53 activity governs this lysosomal exocytosis, we examined how manipulating p53 affects LysoTracker puncta. Activating p53 with Nutlin-3 decreased LysoTracker puncta, while p53 knockdown increased them (**Fig. 4A**), indicating that p53 activation controls lysosomal trafficking and/or acidification. Interestingly, p53 activation with Nutlin-3 combined with HBSS starvation did not further increase STICE unless chloroquine was also added (**Fig. 4G**). Together, these data suggest that starvation drives lysosomal exocytosis specifically in p53-positive cells to trigger extrusion, and that these signalling lysosomes are predominantly non-acidified (**Fig. 4H**). Neither increasing lysosomal deacidification alone (with chloroquine or bafilomycinA1) nor activating p53 alone (with Nutlin-3) was sufficient to increase STICE during starvation; only their combination was, suggesting that these two arms of the pathway synergise rather than act redundantly. In this way, glutamine/glucose starvation induces conventional autophagy in undamaged background cells, but in p53-marked, damaged cells it instead triggers extrusion via lysosomal exocytosis (**Fig. 4H**). It is possible that exocytosis of acidified lysosomes also contributes to degradation of matrix and cell-cell adhesion, though our data suggest that a substantial proportion of these lysosomes are deacidified.

Because STICE selectively removes damaged cells (**Fig. 2**), we tested if starvation might improve overall monolayer fitness by eliminating this defective subpopulation. We found that two hours of nutrient depletion reduced the proportion of γ-H2AX-and p53-positive cells in the monolayer (**Extended Data Fig. 3**), prompting us to test whether the resulting monolayer was more resistant to damage. The remaining cells from starved versus fed monolayers were far more resistant to apoptosis from induction with UV-C^3^ or etoposide^39^ (**Fig. 5A**), as measured by active caspase-3 immunostaining (**Fig. 5B**). Because autophagy can improve survival by recycling damaged components, we tested whether this enhanced survival requires autophagy. However, blocking autophagic flux with chloroquine, did not prevent the survival benefit (**Fig. 5C**). Instead, survival depends on glucose and glutamine depletion, the same nutrients that regulate STICE (**Fig. 5D**). Following starvation, fewer remaining cells were in the cell cycle, as measured by Fluorescent Ubiquitination-based Cell Cycle Indicator (FUCCI)^40^ (**Fig. 5E**). Thus, remaining cells may also be protected from apoptosis by inhibiting cell-cycle re-entry^41^, allowing damage repair. Together, these results suggest that starvation-induced cell-cycle modulation, combined with the selective removal of damaged cells, contributes to the enhanced apoptotic resistance of the monolayer.

**Fig 5:**
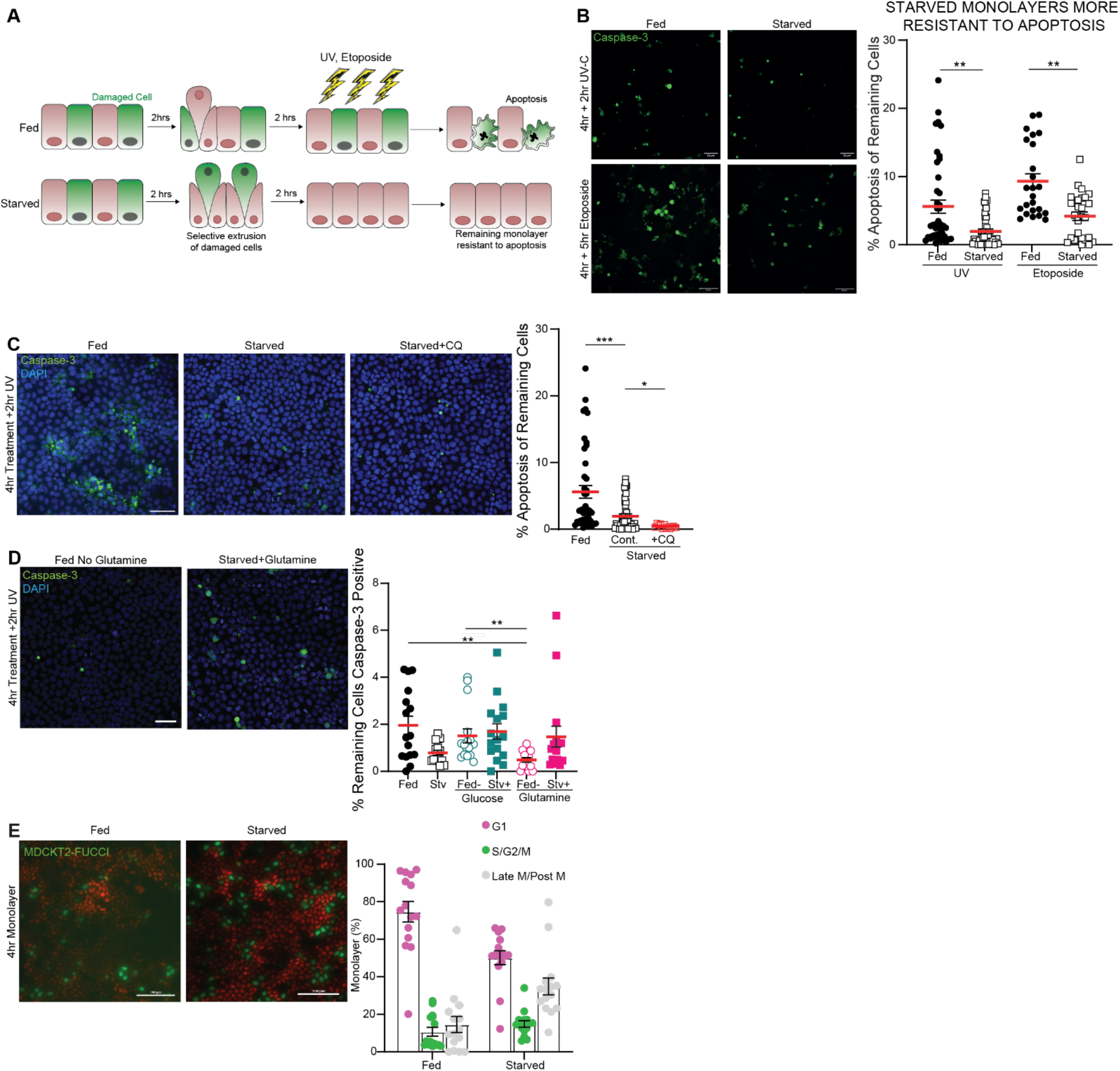
STICE confers enhanced apoptosis resistance. **A**, Schematic showing how starvation may selectively removes damaged cells by extrusion, leaving a monolayer more resistant to apoptosis. **B-D**, MDCKTII cells incubated in DMEM (Fed), HBSS (Starved) (**B**), starved with 50 µM chloroquine (CQ) (**C**), Fed/Starved with glucose or glutamine withdrawn or supplemented (**D**) for 4 h, then treated with UV-C for 2 min followed by a further 2 h incubation or with 50 µM etoposide for 5 h (**B**), to induce apoptosis. Representative confocal images of remaining monolayer (left). Scale bar, 50 µm. Percentage of caspase-3-positive cells remaining in the monolayer (right). **B**, n=40 fields of view (4 biological replicates); p**≤0.01 from Kruskal-Wallis test with Dunn’s multiple-comparisons test. **C**, n=18 fields of view (3 biological replicates); p*≤0.05, p***≤0.001 from Kruskal-Wallis test with Dunn’s multiple-comparisons test. **D**, n=16 fields of view (3 biological replicates); p**≤0.01 from Kruskal-Wallis test with Dunn’s multiple-comparisons test. **E**, Representative live widefield images of FUCCI-MDCKTII cells fed or starved for 4 h (left). Scale bar, 100 µm. Cell-cycle distribution at 4h in fed and starved (right). n=15 fields of view (3 biological replicates).

## Discussion

Here, we show that starvation triggers a regulated wave of extrusion that selectively eliminates sub-fit, DNA-damaged cells or transformed cells. We propose that STICE represents a tissue-level analogue of autophagy: whereas autophagy improves individual cell fitness by digesting damaged components, STICE improves monolayer fitness by eliminating damaged cells (**Fig. 5A**). Extrusion in response to nutrient deprivation also occurs in the sea anemone *Nematostella vectensis*^42^, suggesting this may be a conserved starvation response; however, the underlying mechanism is distinct, driving whole-organism shrinkage rather than the selective removal of defective cells from an otherwise healthy monolayer. While STICE shares elements of the autophagy signalling pathway, it uses some autophagy components to instead trigger lysosomal exocytosis and signal extrusion, rather than to recycle cell contents. Several non-exclusive mechanisms could prime damaged cells for extrusion under starved conditions: 1) release of lysosomal content could reduce cell volume, making the cell more likely to extrude within a crowded environment^10^; 2) released lysosomal hydrolases could degrade surrounding cellular adhesions and promote detachment^37^; 3) or the specifically deacidified lysosomes we noted could act as signalling vesicles that trigger neighbouring cells to extrude the damaged cell^43^. Distinguishing between these mechanisms and identifying what drives them, will require further experimental work.

One clear possibility is that STICE uses a form of conjugation of ATG8 to single membranes (CASM), a non-canonical, ATG16L1-dependent pathway in which LC3 is conjugated directly onto single membranes, including lysosomes, independently of the upstream ULK1-initiated autophagosome formation machinery^44^, consistent with STICE requiring ATG5 but not ULK1 (**Fig. 3B-C**). CASM is generally considered a stress-response pathway that redirects membranes toward faster degradation or secretion. Given that lysosomes in STICE cells are more basic, this elevated pH could itself activate CASM, redirecting defective lysosomes toward exocytosis and out of the cell. Yet CASM alone cannot fully explain this mechanism, which clearly also requires p53 upregulation, potentially redirecting these cells to traffic lysosomes to the plasma membrane. Further studies will be needed to define this mechanism precisely, as it may offer a therapeutic index for targeting cancerous or transformed cells for extrusion without affecting healthy cells.

Our data show that KRas-transformed cells, as well as DNA-damaged cells are selectively eliminated by STICE. Notably, many chemo-resistant cancers also upregulate lysosomal exocytosis to ameliorate the acidosis that stems from the Warburg effect ^45^. Interestingly, fasting-mimicking diets or short-term fasting can improve chemotherapy outcomes and reduce their side-effects ^46–48^. Although it is unclear whether STICE operates under these fasting conditions in humans, if activated it could in principle contribute to this clinical benefit by selectively extruding precancerous or damaged cells while preserving normal tissue. However, most fasting regimes have been developed with autophagy in mind; fasting conditions that specifically induce STICE may therefore require further consideration and development, given that gluconeogenesis buffers glucose levels in the body. Establishing methods to lower glutamine levels in the body may also be challenging. Establishing whether STICE occurs through fasting studies in mice and humans will be important for determining whether fasting benefits tissue through STICE, or whether the benefit prior to chemotherapy is simply due to reduced drug uptake in healthy cells from reduced metabolism and proliferation. If STICE does contribute to this fasting benefit, further identifying the mechanism that regulate it may inform strategies to enhance chemotherapy outcomes.

## Materials & Methods

### Cell Culture

MDCKII cells (European Collection of Authenticated Cell Cultures (ECACC), catalogue number 00062107, lot 19G037) and 16HBE14o (Merck, SCC150) were grown in respective growth medium (**Supplementary Table 1**) supplemented with 10% foetal bovine serum (FBS; 10270106) and 1% penicillin/streptomycin (P/S; 15070063) at 37°C in 5% CO_2_. For all experiments, cells were seeded in growth medium and cultured until confluent. All cell lines tested negative for mycoplasma contamination on periodic testing (Sartorius, 20-700-20).

**Table 1:**
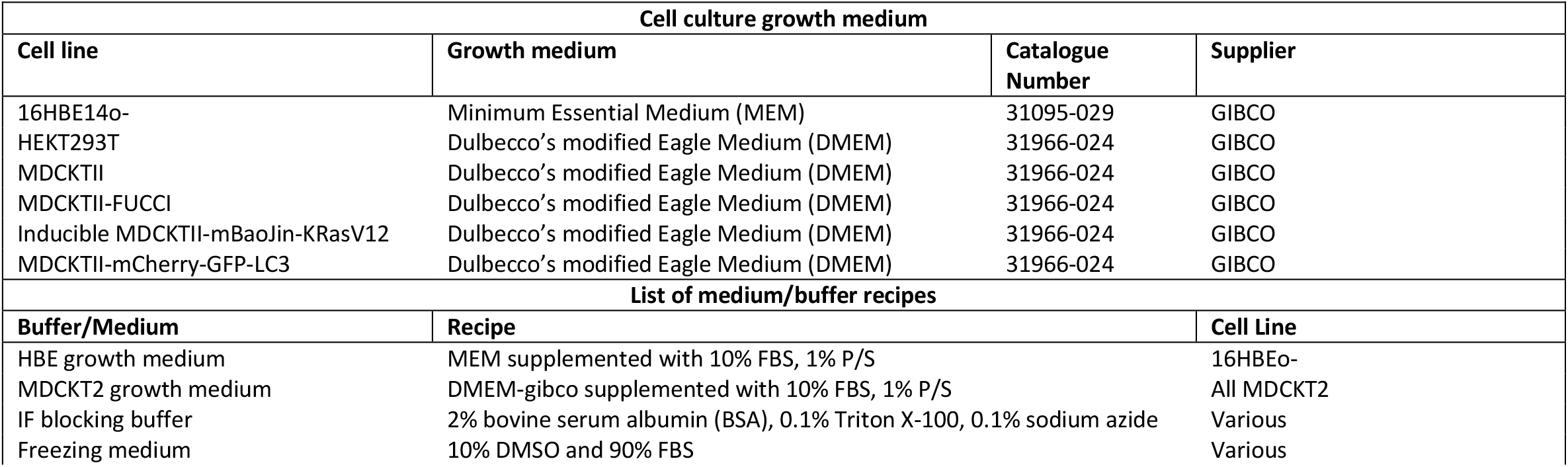
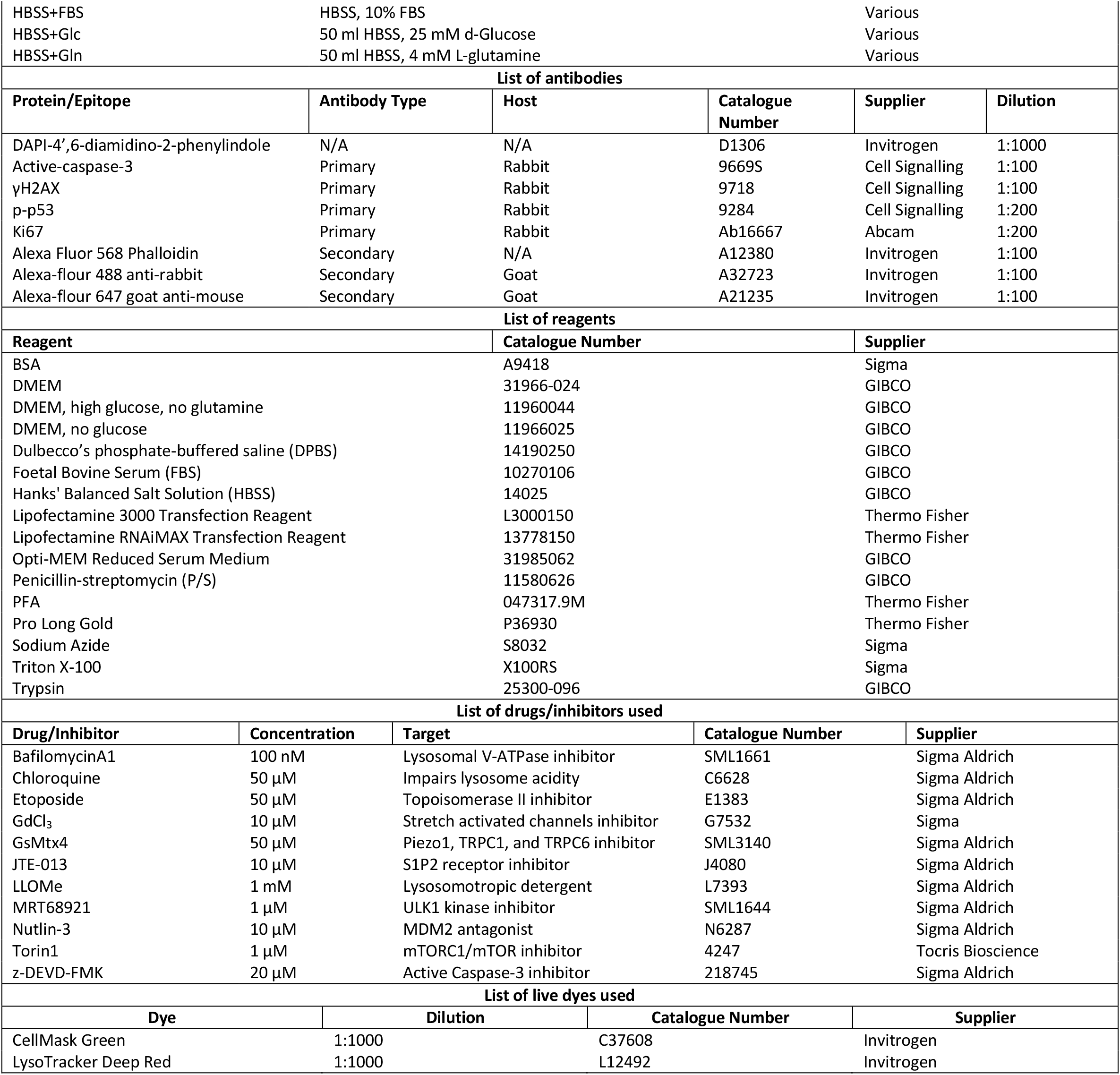
List of reagents and chemicals.

| Cell culture growth medium |  |  |  |
| --- | --- | --- | --- |
| Cell line | Growth medium | Catalogue Number | Supplier |
| 16HBE14o- | Minimum Essential Medium (MEM) | 31095-029 | GIBCO |
| HEKT293T | Dulbecco's modified Eagle Medium (DMEM) | 31966-024 | GIBCO |
| MDCKTII | Dulbecco's modified Eagle Medium (DMEM) | 31966-024 | GIBCO |
| MDCKTII-FUCCI | Dulbecco's modified Eagle Medium (DMEM) | 31966-024 | GIBCO |
| Inducible MDCKTII-mBaoJin-KRasV12 | Dulbecco's modified Eagle Medium (DMEM) | 31966-024 | GIBCO |
| MDCKTII-mCherry-GFP-LC3 | Dulbecco's modified Eagle Medium (DMEM) | 31966-024 | GIBCO |
| List of medium/buffer recipes |  |  |  |
| Buffer/Medium | Recipe | Cell Line |  |
| HBE growth medium | MEM supplemented with 10% FBS, 1% P/S | 16HBEo- |  |
| MDCKT2 growth medium | DMEM-gibco supplemented with 10% FBS, 1% P/S | All MDCKT2 |  |
| IF blocking buffer | 2% bovine serum albumin (BSA), 0.1% Triton X-100, 0.1% sodium azide | Various |  |
| Freezing medium | 10% DMSO and 90% FBS | Various |  |

|  |  |  |  |  |  |
| --- | --- | --- | --- | --- | --- |
| HBSS+FBS | HBSS, 10% FBS |  |  | Various |  |
| HBSS+Glc | 50 ml HBSS, 25 mM d-Glucose |  |  | Various |  |
| HBSS+Gln | 50 ml HBSS, 4 mM L-glutamine |  |  | Various |  |
| List of antibodies |  |  |  |  |  |
| Protein/Epitope | Antibody Type | Host | Catalogue Number | Supplier | Dilution |
| DAPI-4',6-diamidino-2-phenylindole | N/A | N/A | D1306 | Invitrogen | 1:1000 |
| Active-caspase-3 | Primary | Rabbit | 9669S | Cell Signalling | 1:100 |
| γH2AX | Primary | Rabbit | 9718 | Cell Signalling | 1:100 |
| p-p53 | Primary | Rabbit | 9284 | Cell Signalling | 1:200 |
| Ki67 | Primary | Rabbit | Ab16667 | Abcam | 1:200 |
| Alexa Fluor 568 Phalloidin | Secondary | N/A | A12380 | Invitrogen | 1:100 |
| Alexa-flour 488 anti-rabbit | Secondary | Goat | A32723 | Invitrogen | 1:100 |
| Alexa-flour 647 goat anti-mouse | Secondary | Goat | A21235 | Invitrogen | 1:100 |
| List of reagents |  |  |  |  |  |
| Reagent |  | Catalogue Number |  | Supplier |  |
| BSA |  | A9418 |  | Sigma |  |
| DMEM |  | 31966-024 |  | GIBCO |  |
| DMEM, high glucose, no glutamine |  | 11960044 |  | GIBCO |  |
| DMEM, no glucose |  | 11966025 |  | GIBCO |  |
| Dulbecco's phosphate-buffered saline (DPBS) |  | 14190250 |  | GIBCO |  |
| Foetal Bovine Serum (FBS) |  | 10270106 |  | GIBCO |  |
| Hanks' Balanced Salt Solution (HBSS) |  | 14025 |  | GIBCO |  |
| Lipofectamine 3000 Transfection Reagent |  | L3000150 |  | Thermo Fisher |  |
| Lipofectamine RNAiMAX Transfection Reagent |  | 13778150 |  | Thermo Fisher |  |
| Opti-MEM Reduced Serum Medium |  | 31985062 |  | GIBCO |  |
| Penicillin-streptomycin (P/S) |  | 11580626 |  | GIBCO |  |
| PFA |  | 047317.9M |  | Thermo Fisher |  |
| Pro Long Gold |  | P36930 |  | Thermo Fisher |  |
| Sodium Azide |  | S8032 |  | Sigma |  |
| Triton X-100 |  | X100RS |  | Sigma |  |
| Trypsin |  | 25300-096 |  | GIBCO |  |
| List of drugs/inhibitors used |  |  |  |  |  |
| Drug/Inhibitor | Concentration | Target | Catalogue Number | Supplier |  |
| BafilomycinA1 | 100 nM | Lysosomal V-ATPase inhibitor | SML1661 | Sigma Aldrich |  |
| Chloroquine | 50 μM | Impairs lysosome acidity | C6628 | Sigma Aldrich |  |
| Etoposide | 50 μM | Topoisomerase II inhibitor | E1383 | Sigma Aldrich |  |
| GdCl<sub>3</sub> | 10 μM | Stretch activated channels inhibitor | G7532 | Sigma |  |
| GsMtx4 | 50 μM | Piezo1, TRPC1, and TRPC6 inhibitor | SML3140 | Sigma Aldrich |  |
| JTE-013 | 10 μM | S1P2 receptor inhibitor | J4080 | Sigma Aldrich |  |
| LLOMe | 1 mM | Lysosomotropic detergent | L7393 | Sigma Aldrich |  |
| MRT68921 | 1 μM | ULK1 kinase inhibitor | SML1644 | Sigma Aldrich |  |
| Nutlin-3 | 10 μM | MDM2 antagonist | N6287 | Sigma Aldrich |  |
| Torin1 | 1 μM | mTORC1/mTOR inhibitor | 4247 | Tocris Bioscience |  |
| z-DEVD-FMK | 20 μM | Active Caspase-3 inhibitor | 218745 | Sigma Aldrich |  |
| List of live dyes used |  |  |  |  |  |
| Dye | Dilution | Catalogue Number | Supplier |  |  |
| CellMask Green | 1:1000 | C37608 | Invitrogen |  |  |
| LysoTracker Deep Red | 1:1000 | L12492 | Invitrogen |  |  |

### Plasmids

Galectin-3-RFP, mCherry-GFP-LC3, pHLARE and LAMP1-tdtomato were all gifts from Jeremy Carlton (King’s College London).

### Generation of mCherry-GFP-LC3 retroviral stables lines

HEK293T cells were transfected with 500 ng pVSV-G, 2 µg gag-pol and 1 µg mCherry-GFP-LC3 using Lipofectamine 3000 (Invitrogen) diluted in Opti-Mem (Invitrogen), and incubated for 6 h before replacement with fresh medium for a further 48 h. Viral supernatant was collected, centrifuged, and filtered through a 0.2 µm filter before addition to MDCKTII cells seeded in 6-well plate. Cells were incubated for 48 h before selection with 1 mg/ml puromycin for 24 h and expansion into a T25 flask.

### Transfection of MDCKII Cells

MDCKII were seeded at 0.2×10^6^ cells per 6-well plate and grown for 24 h to ∼60% confluence. Cells were transfected with Galectin-3-RFP, pHLARE, or LAMP1-tdtomato using Lipofectamine 3000 (Invitrogen) according to the manufacturer’s protocol for 18 h. Transfection medium was then removed, cells were rinsed twice with PBS and incubated in fresh medium for 24 h before live imaging.

### Harvesting media and re-plating extruded cells

Culture media was collected into a 15 mL tube for secretome analysis. For extruding cells, the remaining monolayer was treated with 500 µL trypsin at room temperature for ∼2 min, with detachment monitored by light microscopy. Trypsinisation was quenched with 2 mL DMEM growth medium and the suspension transferred into a 15 mL tube and centrifuged at 1,200 rpm for 5 min at 4 °C to pellet extruded cells. For re-plating experiments, cells were counted, reseeded at the matched densities on glass coverslips, and grown for 24 h before fixation for immunofluorescence.

### Induction of apoptosis (survival assay)

Confluent MDCKII monolayers were pre-treated with fed (DMEM) and starvation (HBSS) for 4 h at 37°C. Culture medium was centrifuged at 1,200 rpm for 5 min, to remove extruded cells and the resultant supernatant returned to the monolayer. Apoptosis was induced by exposure to UV-C (254 nm) irradiation for 2 min using a handheld UV-C lamp in a UV-safety enclosure, or by addition of 50 µM etoposide for a further 5 h incubation. Cells were fixed and immunostained 2 h after UV-C treatment (or after the 5 h etoposide incubation) for confocal microscopic analysis.

### Immunostaining

Cells were fixed with 4% formaldehyde in PBS for 20 min at room temperature, rinsed three times in PBS, permeabilised for 5 min in PBS containing 0.5% Triton X-100, incubated with primary antibodies (diluted in PBS + 1% BSA; see **Supplementary Table 1** for antibodies and dilutions) overnight at 4°C in a humidified chamber. Following three rinses in PBS, cells were incubated with secondary antibodies (**Supplementary Table 1**) for 2 h at room temperature. Nuclei were counterstained with DAPI (1:1000) for 10 min, washed, and mounted in ProLong Gold antifade mountant, curing for 24 h before imaging.

#### Secretomics

##### Sample Preparation

Media samples were sterile filtered through a 0.45μm PVDF filter (Millex – HV; Merck) prior to secreted proteins being concentrated by spinning through a 3,000 PES molecular weight cut off filter (Vivaspin 500). Samples were then prepared for digestion via S-trap as per manufacturer’s protocol. Briefly, SDS and TEAB were added to the samples to a final concentration of 5% (w/v) and 50mM respectively. Samples were then reduced using 5mM DTT at room temperature for 30 minutes, followed by alkylation with15mM iodoacetamide in the dark at room temperature for 30 minutes. Samples were acidified to pH 1 using phosphoric acid and protein was precipitated through the addition of approximately 6 volumes of 90% methanol, 100mM TEAB. The sample was then loaded onto an S-Trap™ Micro spin column (Protifi), washed 4x with 90% methanol 100mM TEAB, and spun through to remove any residual wash buffer. Trypsin was added at a 1:10 (w/w) ratio in 50mM TEAB and incubated at 37°C overnight. Digested peptides were eluted using 3x elution steps: 50mM TEAB, 0.2% formic acid and 50% acetonitrile; prior to being snap frozen and dried to completion (Labconco CentriVap). Peptides were reconstituted in 0.1% formic acid and desalted using Evotip Pure per Manufacturer’s instructions. Peptides were eluted using 0.1% formic acid in 40% acetonitrile, snap frozen and dried (Labconco Centrivap). Peptides were reconstituted in 0.1% formic acid and peptide concentration was estimated by nanodrop (A280).

##### Mass Spectrometry

Peptides were analysed by LC-MS/MS using a Neo Vanquish (Thermo Scientific) coupled to an Exploris 480 (Thermo Scientific) using a 50 cm, 75 µm I.D. EASY-Spray Pepmap column at a flow rate of 250 nL/min operated in trap and elute mode. Separation was achieved using a 55 minute linear gradient from 2% buffer B to 40% buffer B (buffer A: 0.1% formic acid in water; buffer B: 0.1% formic acid, 80% acetonitrile). The instrument was operated in data-independent acquisition mode. The MS1 scan was performed at a 60,000 resolution, scan range of 380-985, maximum injection time set to 25ms and AGC target set as 300%, and data type set to ‘Profile’. DIA scans were performed in the Orbitrap with a resolution of 30,000, scan range 145-1450 m/z, isolation window size of 20 m/z, window overlap of 1 m/z, an AGC target of 300%, 30% HCD fragmentation, and a maximum injection time of 50ms.

##### Data Analysis

Raw files were analysed using DIA-NN^49^ **version 2.2.0**. All searches were performed in library-free mode with “Deep learning-based spectra, RTs and IMs prediction” enabled, searching against the UniProtKB *Canis lupus familiaris* database (retrieved 27/11/25) including contaminants. Enzyme was set to Trypsin/P and missed cleavages was set to 1. N-terminal methionine excision was selected as a variable modification, and cysteine carbamidomethylation was selected as a fixed modification. Match between runs was turned on. All other settings were left as default. The resulting protein groups table was further analysed using Perseus version^50^ **v2.1.5.0** and R. Protein intensity values were normalised by log(2) transformation and filtered for proteins that were 100% valid in at least one group. Missing values were imputed using a normal distribution (width 0.3, downshift 1.8). For identification of lysosomal proteins, proteins were annotated with GOCC.

##### Live-cell dye staining and imaging

Each dye was diluted in DMEM, enough for all wells, according to their concentration/dilution in **Supplementary Table 1**. Cells were washed 2X PBS, dyed added and incubated for 30 min in the dark. Once stained, cells were washed a further 2X PBS, prior to imaging.

##### siRNA knockdown

SMARTpool siRNAs (Horizon Discovery, L-004374-00-0010 for ATG5; D-001810-01-20 for non-targeting control) and a single-sequence (Horizon Discovery, J-003329-17-0005 for TP53) were resuspended in DNase/RNase-free water to a 100 μM stock. MDCKTII cells were seeded at 0.2×10^6^ cells per well in a 6-well plate and grown for 24 h to ∼ 60% confluence, then transfected with 1 µM siRNA using RNAiMAX (Invitrogen) for 24 h before replacement with fresh DMEM for 48 h. Cells were then given fresh treatments (DMEM or HBSS) and imaged to monitor extrusion. Extrusions were quantified as described below, and knockdown efficiency was confirmed by western blot.

##### Widefield microscopy

Time-lapse phase-contrast and fluorescence images were captured on a Nikon Eclipse Ti2 system using a Plan Fluor 20x Ph1 DLL NA=0.50 objective, a Photometrics Iris 15 16-bit camera and a CoolLED pE-4000 illumination source, driven by NIS-Elements software (**Nikon, v.5.30.02**).

##### Spinning-disk microscopy

Images were captured on a Nikon Eclipse Ti2 system using Plan Fluor 20x, 40x Plan Fluor water or 60X, 100X Plan Fluor 1.40 oil objectives with an iXon 888 Andor 16-bit camera, a Yokogawa CSU-W1 confocal spinning disk unit and Toptica photonics laser driven by NIS-Elements software (**Nikon, v.5.21.03**). Phalloidin- and Hoechst-stained cells were quantified for extrusion rate (per 1,000 or 10,000 cells, as indicated in figure legends) using NIS-Elements Software.

##### Extrusion quantification

Extrusion events were quantified from time-lapse phase-contrast videos of MDCKTII monolayers by identifying cells eliminated from the monolayer by classical extrusion. Extruding cells were distinguished from dividing cells by absence of two daughter cells reincorporating into the monolayer. Cells that rounded up, divided and reincorporated were scored as mitoses. Extruding cells instead rounded up, became phase-white, and either detached and floated free or remain transiently attached at the monolayer surface. Cells already extruded at the start of filming were excluded from quantification.

##### TrackMate analysis of LysoTracker

TrackMate plugin^51^ of ImageJ was used. ROIs were manually drawn around the plasma membrane, stained with CellMask, of representative single cells in the monolayer and movies were cropped to remove the latter few time points to minimize when the dye faded and any drift in imaging. The channels were split to focus on the LysoTracker channel, which was selected for TrackMate. Puncta were thresholded based on quality, diameter, linking distance, and max frame gap. LoG detector was used to detect puncta, tracks were made with “simple LAP tracker”. Tracks were then analysed with MatLab.

## Supporting information

Phase microscopy timelapse (h:min) of MDCKTII cells treated with HBSS (starvation). Images were taken every 10 min.

Phase microscopy timelapse (h:min) of MDCKTII cells treated with complete medium (Fed). Images were taken every 10 min.

Confocal fluorescence microscopy timelapse (h:min:sec) of MDCKTII-mCherry-GFP-LC3 cell extruding in HBSS (starvation), showing STICE cells unable to f

Confocal fluorescence microscopy timelapse (h:min:sec) of MDCKTII-mCherry-GFP-LC3 non-extruding in HBSS (starvation), showing monolayer cell able to f

LysoTracker (magenta) and CellMask (green) timelapse (h:min:sec) overlayed with LysoTracker tracks from TrackMate of a STICE cell pre-extrusion, showi

LysoTracker (magenta) and CellMask (green) timelapse (h:min:sec) overlayed with LysoTracker tracks from TrackMate of a starved monolayer cell, showing

Fluorescence microscopy timelapse (h:min:sec) of MDCKTII-LAMP1-tdtomato (red) pre-treated for 1h starvation (HBSS), showing lysosomal traffic to the c

Fluorescence microscopy timelapse (h:min) of MDCKTII-LAMP1-tdtomato (red) cells extruding. Cell mask (green). Same field of view as Supplementary Vide

Widefield fluorescence and phase microscopy timelapse (h:min:sec) of MDCKTII-pHLARE treated in HBSS, including both extruding and non-extruding monola

## Statistics and reproducibility

All experiments were repeated independently on at least three separate days to capture biological variability. Data were analysed in GraphPad Prism **v.10.6.0**. Normality was assessed using Shapiro-Wilk test. Statistical significance was determined using unpaired or ratio-paired *t*-tests (all *t*-tests were performed with two-tailed analysis), two-way ANOVA with Tukey’s or Šidák’s correction, or one-way ANOVA with Dunnett’s or Welch’s correction, as specified in individual figure legends. To minimise selection bias, imaging fields were chosen at random within the middle/crowded regions of glass coverslips for fixed-cell quantification, or the centre of an 8-well imaging dish (Ibidi, 80806) for live-cell imaging experiments. Low-density epithelia (fewer than 2,000 cells per field) were not considered sufficiently crowded to elicit extrusion under fed conditions. Graphs were generated using GraphPad Prism **v.10.6.0**; figure layouts and schematic models were created in Adobe Illustrator **v.28.7.1**.

**Extended Data Figure 1:**
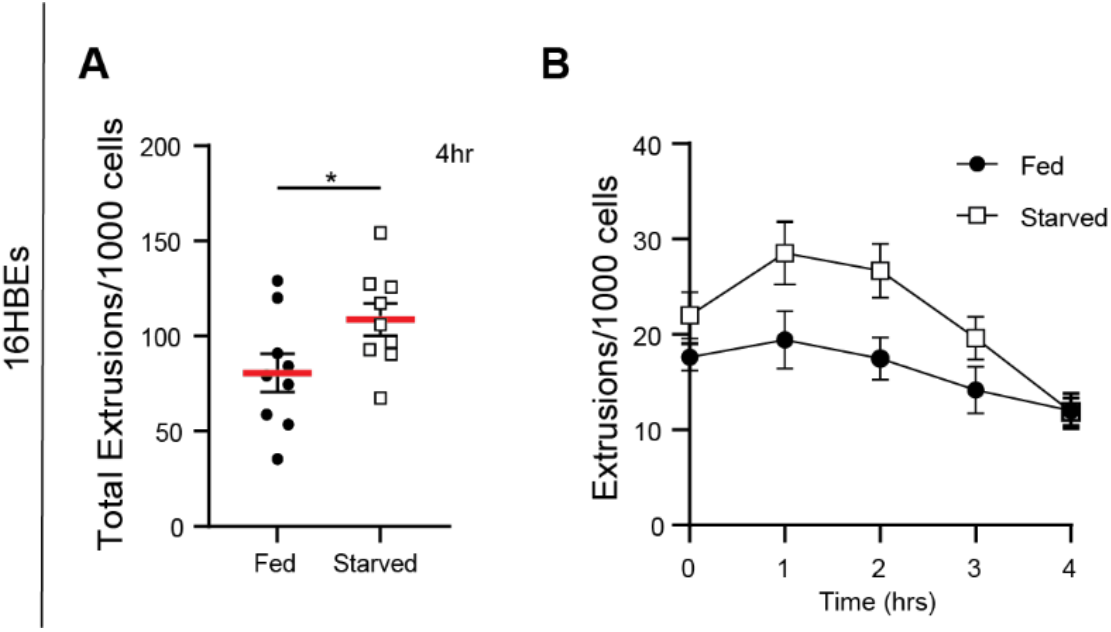
STICE is a conserved process. 16HBE cells cultured in normal growth medium (Fed) or HBSS (Starved), imaged every 10 min for 4 h. **A**, Normalised extrusions per 1,000 cells over the 4 h time course. n=9 fields of view (3 biological replicates). Statistical analysis was performed using an unpaired t-test; p*≤0.05. **B**, Time course of total extrusions (normalised per 1,000 cells) scored hourly. n=9 fields of view (3 biological replicates).

**Extended Data Figure 2:**
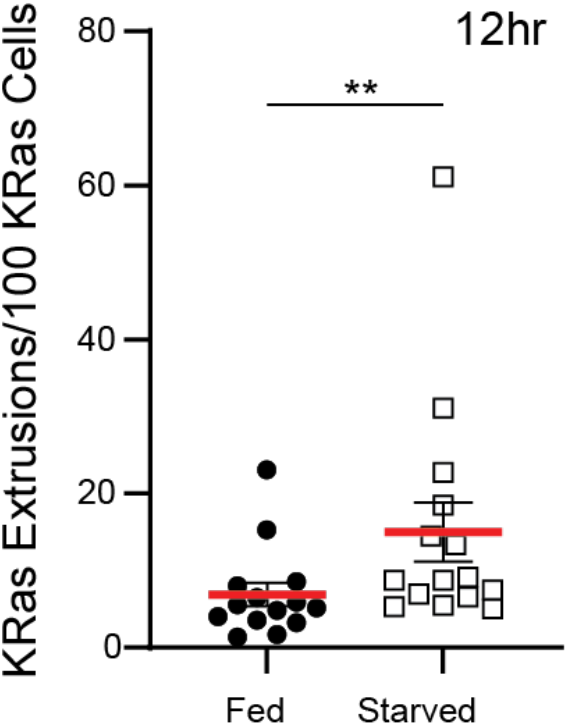
Starvation induces extrusion of KRasV12-transformed cells. Wild-type and doxycycline-inducible GFP-KRasV12 MDCKTII cells combined 50:1, induced 24 h prior to imaging, incubated in DMEM (Fed) or HBSS (Starved), and imaged for 12 h at 10 min intervals. Total KRasV12-positive extrusions (normalised per 100 KRasV12 cells) over 12 h. n=3 fields of view (5 biological replicates). Statistical analysis was performed using a Mann-Whitney t-test; p**≤0.01.

**Extended Data Figure 3:**
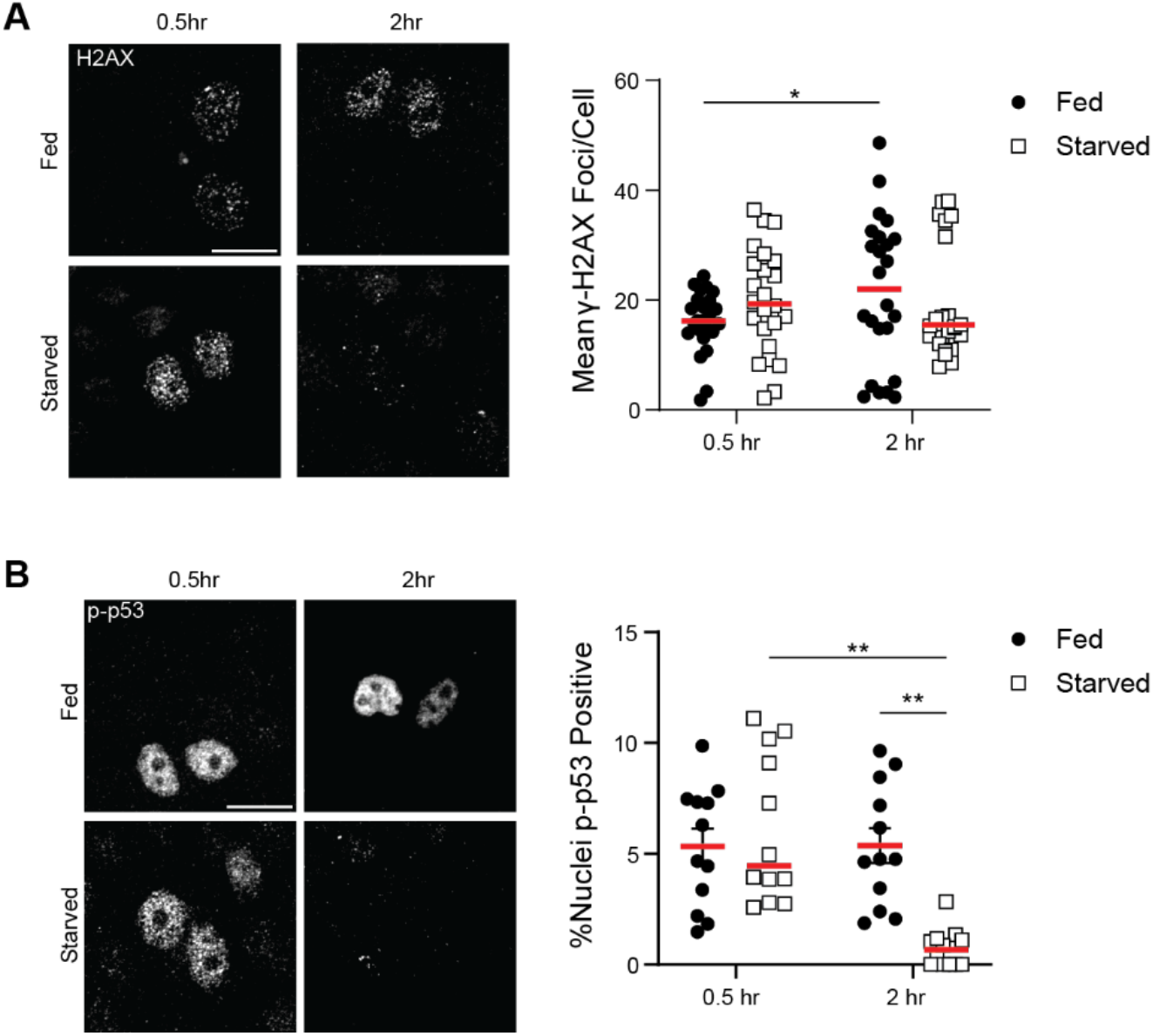
Starvation reduces DNA damage within the monolayer. MDCKTII cells incubated in either DMEM (fed) or HBSS (starved) for 0.5 or 2 h, fixed and stained for γ-H2AX (**A**) and p-p53 (**B**). Representative confocal images (left). Scale bar, 10 µm. γ-H2AX foci per cell (**A**, right) and percentage of p-p53-positive nuclei (**B**, right). n=25 fields of view (5 biological replicates). Statistical analysis was performed using repeated-measures ANOVA with Šidák’s multiple-comparisons test (**A**) or one-way ANOVA (**B**); p*≤0.05, p**≤0.01.

**Supplementary Video 1:**

**Supplementary Video 2:**

**Supplementary Video 3:**

Confocal fluorescence microscopy timelapse (h:min:sec) of MDCKTII-mCherry-GFP-LC3 cell extruding in HBSS (starvation), showing STICE cells unable to flux their LC3. Images were taken every 1 min. Scale bar 10 µm.

**Supplementary Video 4:**

Confocal fluorescence microscopy timelapse (h:min:sec) of MDCKTII-mCherry-GFP-LC3 non-extruding in HBSS (starvation), showing monolayer cell able to flux their LC3. Images were taken every 1 min. Scale bar 10 µm.

**Supplementary Video 5:**

LysoTracker (magenta) and CellMask (green) timelapse (h:min:sec) overlayed with LysoTracker tracks from TrackMate of a STICE cell pre-extrusion, showing a LysoTracker movement to the plasma membrane. Images were taken every 15 s. Scale bar 10 µm.

**Supplementary Video 6:**

LysoTracker (magenta) and CellMask (green) timelapse (h:min:sec) overlayed with LysoTracker tracks from TrackMate of a starved monolayer cell, showing a static LysoTracker signal. Images were taken every 15 s. Scale bar 10 µm.

**Supplementary Video 7:**

Fluorescence microscopy timelapse (h:min:sec) of MDCKTII-LAMP1-tdtomato (red) pre-treated for 1h starvation (HBSS), showing lysosomal traffic to the cell membrane and membrane buckling in pre-extruding cells, not seen in non-extruding cells. Images were taken every 15s. Scale bar 10 µm.

**Supplementary Video 8:**

Fluorescence microscopy timelapse (h:min) of MDCKTII-LAMP1-tdtomato (red) cells extruding. Cell mask (green). Same field of view as **Supplementary Video 7**, 2h into starvation treatment. Images were taken every 5 min. Scale bar 10 µm.

**Supplementary Video 9:**

Widefield fluorescence and phase microscopy timelapse (h:min:sec) of MDCKTII-pHLARE treated in HBSS, including both extruding and non-extruding monolayer cells. Images were taken every 10 min. Scale bar 25 µm.

## Acknowledgements

We thank Jeremy Carlton for his advice throughout this project and for providing all plasmids, Alexandros Sanchez Vassopoulos and Gabriela Vino Flores for the generation of the inducible MDCKTII-mBaoJin-KRasV12 line, and members of the Rosenblatt laboratory for help with experimental design and figure layouts feedback.

## Author Contributions

J.R., L.G. and D.C.B. conceived the study and designed the experiments. L.G. performed all experiments and data analyses. L.G. and J.R. interpreted the data and wrote the manuscript. K.K. interpreted and assisted with the TrackMate analysis. T.A., and M.V. assisted with secretomics data acquisition. T.A., M.V., and J.F., interpreted secretomics data. All authors edited and approved the final manuscript.

## Data and Materials Availability

All data are available in the main text or the extended data. The source material link will be provided upon request.

